# Activin A promotes bone fracture repair and acts through a novel myofibroblastic cell population in callus

**DOI:** 10.1101/2022.10.05.510962

**Authors:** Lutian Yao, Leilei Zhong, Yulong Wei, Tao Gui, Luqiang Wang, Jaimo Ahn, Joel Boerckel, Danielle Rux, Christina Mundy, Ling Qin, Maurizio Pacifici

## Abstract

Insufficient bone fracture repair represents a significant clinical burden, and identification of novel therapeutics enhancing repair would have substantial clinical and societal impact. Activin A is a TGF-β protein superfamily member known to stimulate ectopic bone formation, but its roles in fracture repair and its therapeutic potentials remain unclear. Using two mouse tibia fracture repair models, here we mapped activin A expression at the tissue and single cell levels, tested its requirement for normal repair and evaluated its ability to enhance repair when provided exogenously. Activin A was minimally expressed in periosteum of intact bones but was markedly upregulated in developing callus soon after fracture. Single cell RNA-sequencing revealed that the activin A-encoding gene *Inhba* marked a unique, highly proliferative progenitor cell (PPC) population with a myofibroblast character which emerged over repair time and lay at the center of a developmental trajectory bifurcation producing cartilage and bone cells within callus. Systemic administration of a neutralizing activin A antibody impaired fracture repair and its endochondral and intramembranous phases, whereas local delivery of recombinant activin A enhanced repair. Activin A delivery also induced SMAD2 phosphorylation in vivo and increased the fraction of αSMA+ myofibroblasts within fracture callus. Gain- and loss-of-function experiments in vitro showed that activin A directly stimulated myofibroblast differentiation, chondrogenesis and osteogenesis in periosteal progenitor cells. Together, these data identify a unique population of *Inhba*-expressing proliferative progenitor cells that give rise to chondrocytes and osteoblasts during fracture healing and establish activin A as a potential new therapeutic tool to enhance it.

**One Sentence Summary:** Deficits in bone fracture repair remain a clinical challenge and the present study provides evidence for the therapeutic potentials of activin A

## INTRODUCTION

Bone is endowed with the ability to heal after fracture and does so effectively in the majority of, but not all, patients (*1, 2*). Given its traumatic nature, a bone fracture triggers local hematoma formation followed by an inflammatory cascade that involves innate and adaptive immune responses (*3*). These initial steps elicit the expansion and migration of the otherwise quiescent mesenchymal stem and progenitor cells within the periosteum that over time undergo differentiation into chondrocytes within the body of callus and osteoblasts at its peripheral ends. Subsequent remodeling of callus routinely leads to nearly scar-less bone healing. Thus, fracture healing replicates, and relies on, many of the processes of endochondral and intramembranous ossification through which the skeleton normally develops and grows prenatally and postnatally (*4*). Recent studies have aimed to clarify the character of the periosteal progenitors and their roles in the fracture repair process. Building on their earlier reports (*5, 6*), Matthews et al. characterized a *αSMA*-expressing slow-cycling, long-term and self-renewing periosteal progenitor cell population that when ablated, impaired healing (*7*). Another report provided evidence of a periosteum-resident durable Mx1+αSMA+ stem cell subpopulation that expressed chemokine CCL receptors needed for effective bone healing (*8*). Despite these and other advances, much remains unclear about the regulation of fracture repair and in particular: what factors set the repair process in motion and in what sequence they act; what distinguishable stages exist along the repair process and how cells move from one to the next; how the progenitors commit to separate differentiation lineages; and importantly for patient care, to what extent the progenitor cell’s repair capacity can be modified, corrected or improved. Unbiased single cell RNA-sequencing (scRNA-seq) approaches have recently been used to characterize mesenchymal cells in marrow, delineating their regulation, functioning and heterogeneity (*9-12*).

Similar approaches applied to the periosteum could elicit similar fundamental insights into the regulation and roles of progenitor cells in bone repair mechanisms, as the present study shows. These discoveries would also hold high clinical significance and importance as such knowledge could be leveraged to mitigate healing deficiencies seen in about 5 to 10% of patients, leading to problematic and costly non-unions that are not fully addressed by current therapeutic strategies (*13, 14*). Thus, new directions and options are needed.

As it has been observed previously (*15*), fracture repair shares multiple phenotypic traits and steps with heterotopic ossification (HO), a pathology in which skeletal tissue forms and accumulates at orthotopic locations. One form of HO is congenital, rare and often fatal and characterizes patients with Fibrodysplasia Ossificans Progressiva (FOP) (*16*), and there is an acquired, common and potentially debilitating form of HO that can occur in individuals after invasive surgeries, physical trauma, deep burns or spinal cord injury (*17-19*). Both congenital and acquired HO most often initiates with local inflammation that is followed by recruitment of progenitor stem cells, skeletogenic cell differentiation and local deposition of cartilage and bone, steps indeed reminiscent of those underlying the fracture repair process. Recent studies from this and other laboratories have revealed that the transforming growth factor-β (TGF-β) superfamily member activin A has a previously unsuspected role in promoting congenital and acquired forms of HO in mice (*20-22*). This protein is well known for its required roles in several fundamental physiologic processes including immune responses, hematopoiesis and wound healing (*23-25*). Mice lacking the activin A-encoding gene, *Inhba*, die at birth (*26*) and changes in activin A availability, levels and function contribute to certain pathological conditions (*23, 25*). Because of these observations, we tested here whether activin A is necessary for fracture repair in mice and whether exogenously provided recombinant protein would promote it, offering a new potential therapeutic strategy. The data in this report provide support for this exciting new direction. In particular, our scRNA-seq data and computational analysis of the periosteum identify a novel myofibroblast-like cell population that quickly expands after fracture, expresses *Inhba* and gives rise to chondrocytes and osteoblasts, promoting bone healing.

## RESULTS

### Levels and distribution of activin A are elevated during fracture repair

To evaluate the participation of activin A in fracture repair, we first asked whether its presence and distribution were altered during healing. To this end, we utilized a closed mouse model (*27*) in which a controlled traumatic fracture is introduced with a blunt guillotine on the right tibia of 2 month-old *WT* mice bearing a pre-inserted intramedullary pin for stabilization used in our previous studies (*28*). Immunofluorescent staining of intact control tibiae (Fig. 1A) showed that activin A was detectable in only a few cells within the cambium layer of periosteum (Fig. 1Aa) and in some bone marrow cells (Fig. 1Ab). Marrow cells normally expressing the activin A-encoding gene *Inhba* included mesenchymal progenitors and inflammatory cells such as granulocytes, based on our previous scRNA-seq analysis (fig. S1) (*29*). Five days after fracture, the number and distribution of activin A-positive cells had dramatically increased (Fig. 1B), and expressing cells now included numerous mesenchymal progenitors within periosteum (Fig. 1Ba) and round-shaped early chondrocytes within the developing soft callus (Fig. 1Bb). Clearly, activin A+ cells become abundant during the early stages of fracture repair.

**Fig. 1.**
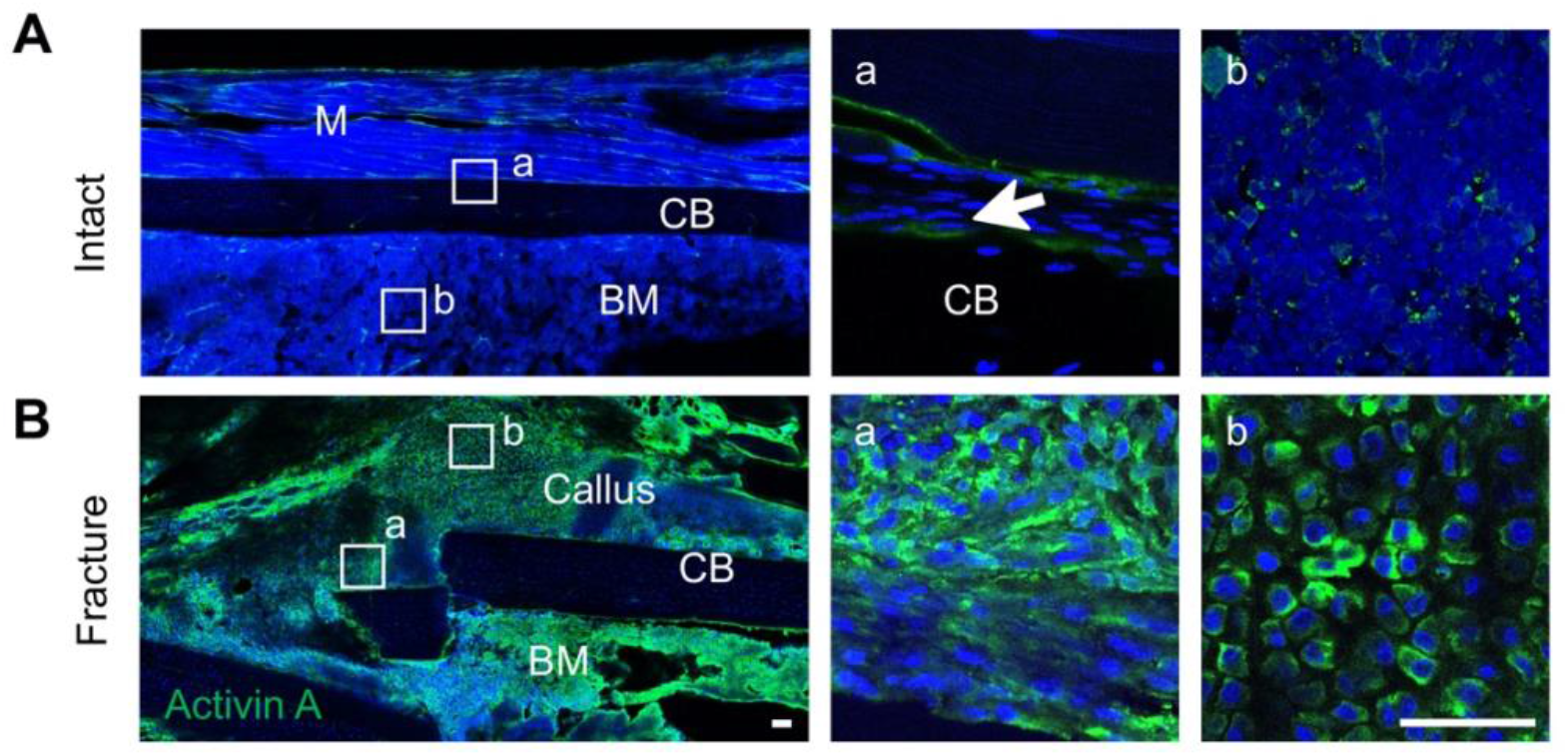
Activin A becomes more abundant during fracture repair. (**A**) Immunofluorescence images of activin A distribution in intact mouse tibia. The outlined periosteum and bone marrow areas on the left panel are shown as enlarged images in a and b, respectively, on the right. (**B**) Immunofluorescence images of activin A distribution in mouse tibial callus at day 5 post fracture. The boxed periosteum and early cartilage areas on the left panel are shown as enlarged images as a and b, respectively, on the right. M: muscle, CB: cortical bone, BM: bone marrow. Scale bar: 50 μm.

### Fracture callus development involves dynamic cell population shifts

Before testing the possible roles of activin A in fracture healing directly, we next carried out a comprehensive and unbiased analysis of cell populations in the fracture repair process and possible links to *Inhba* expression, using scRNA-seq. To trace, track and sort the cell populations involved, we carried out fracture repair in *Col2-Cre;Gt(Rosa)26 tdTomato* (*Col2/Td*) reporter mice. We selected this mouse model because as our previous study indicated, Td+ cells in these mice contribute to fracture callus (*30*). To establish the effectiveness of this transgenic approach to capture the overall cell populations involved in fracture repair, tibiae from operated 2 month-old *Col2/Td* mice were harvested at 5, 10, 15 and 30 days after fracture and processed for spatiotemporal delineation of Td+ cells (fig. S2). In intact tibiae, the cortical bone surface was covered by a thin layer of periosteum consisting of Td+ cells (fig. S2, Aa and Ba). At day 5 post fracture, Td+ cells had greatly expanded in number to form the thickened periosteum (fig. S2Ab, Bb). By day 10 when cartilage reached its peak soft callus development as shown by Safranin O staining, all chondrocytes were Td+ as were nearby fibrotic cells (fig. S2, Ac and Bc). By day 15 when most cartilage was undergoing endochondral ossification, Td+ cells now constituted the majority of osteoblasts and osteocytes in the callus (fig. S2, Ad, Bd). In the remodeling bone present by day 30, the new periosteum at the edge of callus consisted of mostly Td+ cells (fig. S2, Ae and Be). CFU-F assays of periosteal cells isolated from intact bones revealed that Td+ cells, but not Td-cells, were able to form cell colonies (fig. S2, C and D), affirming that the Col2/Td approach captured the overall mesenchymal cell populations taking part in fracture healing.

Having established the effectiveness of the *Col2/Td* approach, we proceeded to isolate periosteal cells from intact tibiae (termed day 0 cells) and from injured tibiae at day 5 and day 10 post fracture and then sorted them for Td+ cells. The percentage of Td+ cells amongst freshly isolated cells increased from 2.8% at day 0 to 3.4% at day 5 and 8.5% at day 10 (fig. S3A). Using 10x Genomics approach, we sequenced 7496, 7535 and 10398 Td+ cells from day 0, 5, and 10 samples respectively, with a median of 3462 genes/cell and 13463 unique molecular identifiers (UMIs)/cells (fig. S3B). We merged the resulting three datasets comprising an overall total of 25429 cells that elicited 16 cell clusters, including 6 clusters of periosteal mesenchymal lineage cells, 4 clusters of hematopoietic cells, 1 cluster of muscle cells, 1 cluster of synovial lining cells, 1 cluster of tendon cells, 1 cluster of endothelial cells (ECs), 1 cluster of neuron cells, and 1 cluster of smooth muscle cells (SMCs) (fig. S3, C and D). *Td* expression was detected in all clusters but was highest in mesenchymal cell lineages (fig. S3E). There was some basal *Td* expression in non-mesenchymal cells and the same was observed in our previous study of bone marrow mesenchymal lineage cells using this approach (*9-11*) and in similar scRNA-seq studies from other groups (*9-11*). Heatmap analysis and violin plots revealed the hierarchy and diverse nature of these cell clusters with distinct gene signatures (fig. S3, D and F). After digitally removing the non-mesenchymal clusters, the recalculated data from the mesenchymal lineage populations elicited 6 distinct clusters (Fig. 2, A and B). Based on lineage specific traits, clusters 3, 4, 5, and 6 represented early osteoblasts (EOB), osteoblasts (OB), chondrocytes (CH) and hypertrophic chondrocytes (HC), respectively (Fig. 2C). Cells in cluster 1 were characterized by several typical stem cell markers such as *Cd34, Ly6a* and *Thy1*, suggesting that they represented mesenchymal progenitor cells (MPCs). Cells constituting the expansive cluster 2 expressed the above stem cell markers at a level lower and were also largely devoid of lineage specific gene markers such as those of EOBs and CHs. Notably, cluster 2 cells exhibited strong expression of genes characteristic of myofibroblasts, including *αSMA* (*Acta2*) (*31*). When the merged datasets (Fig. 2A) were separated by time point, it became clear that cluster 2 cells markedly increased in number early from day 0 to day 5, whereas chondrocytes (cluster 5), hypertrophic chondrocytes (cluster 6) and osteoblasts (clusters 3 and 4) had expanded by day 10 (Fig. 2B and fig. S4). Interestingly, computational cell cycle analysis revealed that cluster 2 contained highly proliferative cells, particularly at the day 5 time point (Fig. 2, D and E). Considering their overall phenotypic traits and behavior, we termed -and refer to-cluster 2 cells as proliferative progenitor cells (PPCs).

**Fig. 2.**
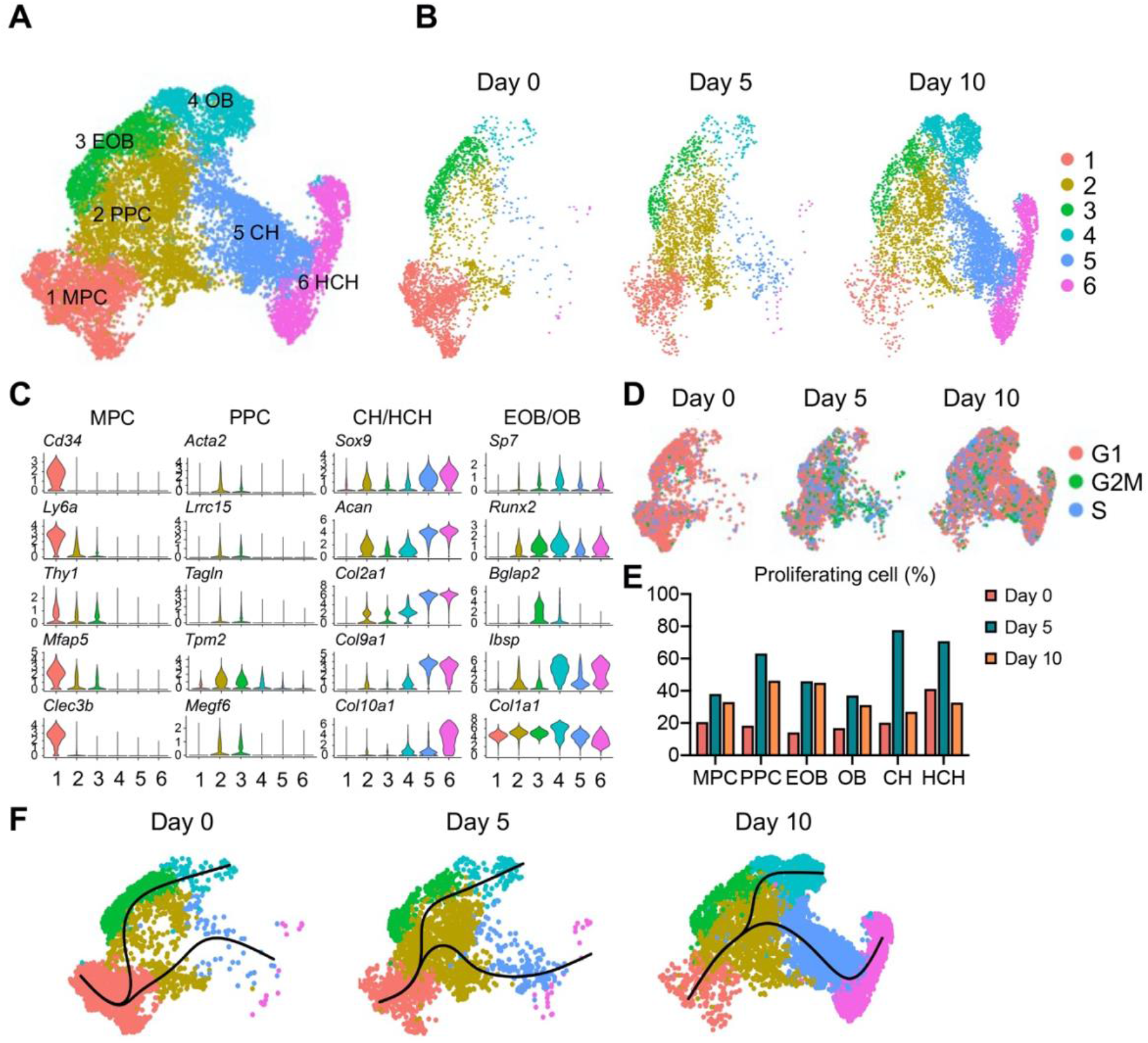
Single cell transcriptomics analyses reveal the identity and developmental trajectories of periosteal mesenchymal populations. (**A**) The UMAP plot of 13040 Td+ mesenchymal lineage cells isolated from tibial periosteum of 2 month-old *Col2/Td* mice. Datasets from cells isolated from intact periosteum (day 0) and fracture site on days 5 and 10 post-surgery were merged and combined into a single plot. (**B**) The UMAP plots of those cells shown at each respective time point. (**C**) Violin plots of cluster-specific makers of mesenchymal lineage cells. MPC: mesenchymal progenitor cell; PPC: proliferative progenitor cell; CH: chondrocyte; HCH: hypertrophic chondrocyte; EOB: early osteoblast; OB: osteoblast. (**D**) Cell cycle phase of periosteal mesenchymal lineage cells at days 0, 5 and 10 as above. (**E**) Percentages of proliferative cells (S/G2/M phase) in each cell cluster at three time points above was computationally quantified. (**F**) Slingshot trajectory plots of periosteum mesenchymal lineage cells at days 0, 5 and 10.

Examination of previously reported markers of periosteal mesenchymal progenitors in our dataset revealed that most of them are ubiquitously expressed in all subpopulations (fig. S5). Some genes are expressed equally among all clusters, such as *Itgb1* (Cd29), *Itgav* (Cd51), *Eng* (Cd105), *Gli1* and *Lepr* (*32-35*), and some are expressed at a higher level in progenitors and osteoblasts than in chondrocytes, such as *Ctsk, Postn, Prrx1* and *Pdgfrb* (*36-39*). One marker, *Cd200* (*34, 35*), is expressed equally in all cells but at a low level in MPCs. Another two markers, *Col2a1* (*30*) and *Sox9* (*40*), are expressed at a much higher level in chondrocytes than the rest of mesenchymal cells, which is consistent with the conventional wisdom that they are cartilage markers. We did not detect the expression of a previously proposed marker *Mx1* (*41*) in our dataset.

RNA velocity delineates cellular differentiation paths and transient phenotypic states from scRNA-seq data (*42*). Applying this approach to our merged datasets above (Fig. 2A), we found that directionality of cell differentiation and diversification started from cluster 1 (MPCs), advanced and transitioned through cluster 2 (PPCs) and ended in cluster 4 (OBs) and cluster 6 (HCH) (fig. S6). Likewise, at all time points, pseudotemporal cell trajectory analysis placed MPC cells (cluster 1) at one end of the developmental trajectory, PPC cells (cluster 2) in a central position, and OBs (cluster 4) and CHs (clusters 5 and 6) at the other two ends (Fig. 2F).

Together, the data strongly indicate that MPCs serve as stem/progenitors and give rise to myofibroblastic PPCs which in turn diverge into chondrocytes and osteoblasts, contributing to soft and hard callus formation.

### PPCs strongly express *Inhba* and possess a myofibroblast character

Given the apparent developmental centrality of the PPC population, we sought to characterize it further by defining their differentially expressed genes (DEGs) compared to those in the other cell clusters, using GO term and KEGG analyses (Fig. 3A). Importantly, the most up-regulated genes in PPCs verified their myofibroblast character. Those genes were closely related to processes and pathways known to be regulated by myofibroblasts such as wound healing, cell adhesion, extracellular matrix organization and contractile actin filament (*31*), and included *Acta2, Tagln and Myl9* (Fig. 3B and fig. S7A). Conversely, the least expressed genes were those related to ossification, mineralization and chondrocyte differentiation, confirming that the PPCs did not possess a terminally differentiated phenotype (Fig. 3, A and B). Of particular relevance to the present study were the findings that (i) the PPCs prominently expressed *Inhba* and did so at a level far higher than MPCs, EOBs and OBs but comparable to CHs (Fig. 3C and fig. S7B) and (ii) *Inhba* expression increased along with the increase in PPC number and their developmental bifurcation into chondrocytes and osteoblasts over time (Fig. 3C and fig. S7B).

**Fig. 3.**
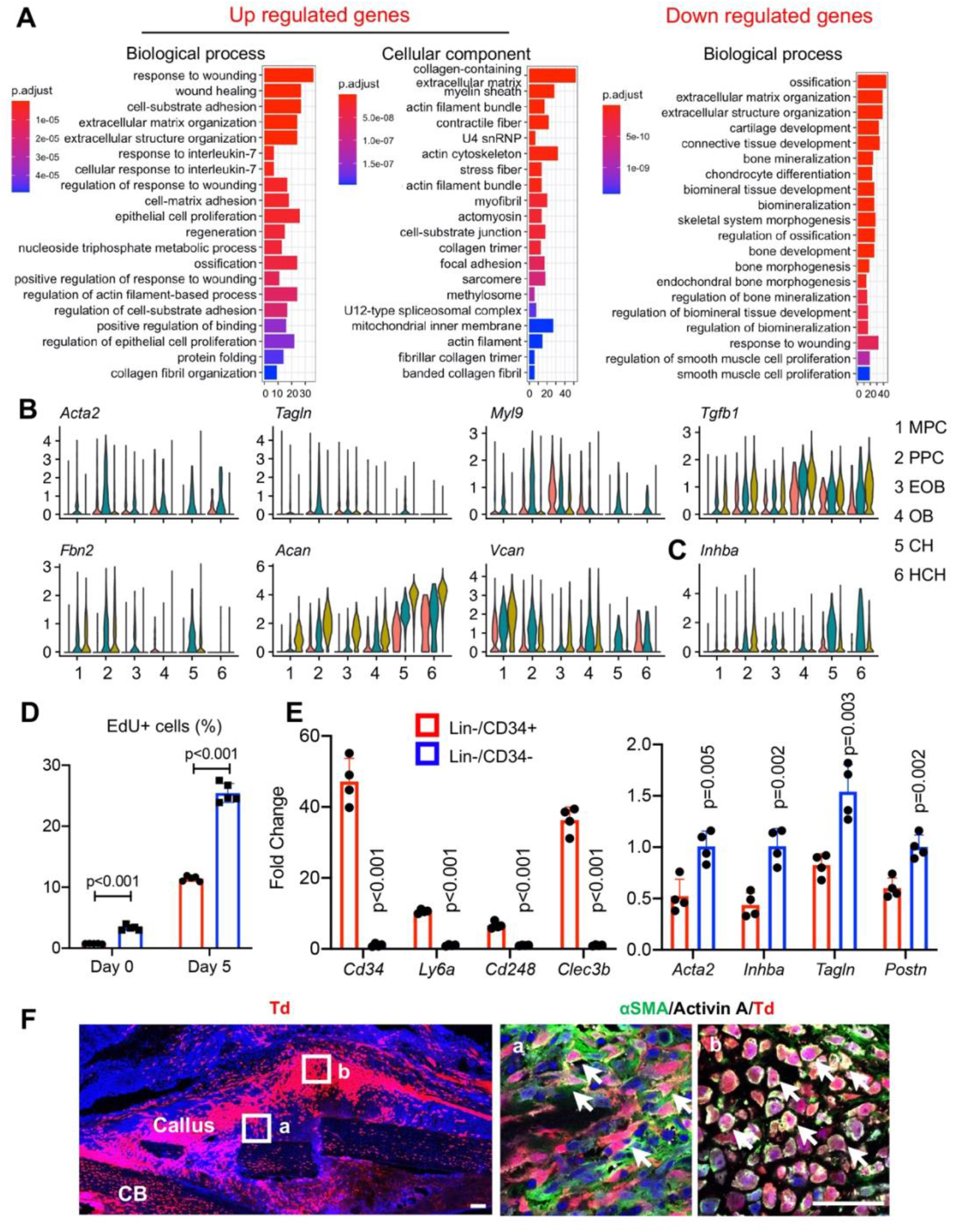
PPCs have a myofibroblast phenotype and express *Inhba*. (**A**) GO term analysis of genes up-regulated or down-regulated in PPC cluster compared to the rest of periosteal mesenchymal cell clusters. (**B**) Violin plots of myofibroblast marker and extracellular matrix gene expression. (**C**) Violin plots of *Inhba* gene expression. (**D**) Calculated percentages of EdU+ cells in MPCs (Lin-CD34+) and PPCs (Lin-CD34-) from mouse periosteum at day 0 before fracture and day 5 post fracture. Mice received an EdU injection at 24 hr before bone harvest. n= 4 mice/group. (**E**) qRT-PCR analyses comparing the expression of stem cell markers, myofibroblast markers, and *Inhba* in MPCs and PPCs at day 5 post fracture. n = 4 mice. (**F**) Immunofluorescence images of αSMA and activin A distribution in mouse callus at day 5 post fracture. Boxed periosteum and early cartilage areas on left panel are shown enlarged as a and b, respectively, on the right. CB: cortical bone. Scale bar: 50 μm.

Together, the data strongly suggest that *Inhba* is a novel myofibroblast cell marker. To further validate the above findings and considerations, we focused on the interval between day 0 to 5 when the PPCs increased the most in number (Fig. 2B) and sorted Cd45-Cd31-Ter119-Cd34+ (Lin-/Cd34+) cells and Cd45-Cd31-Ter119-Cd34- (Lin-/Cd34-) cells to further delineate their character. Based on scRNA-seq data, the day 0 cells mainly represented MPCs (Lin-/Cd34+) with relatively few EOBs and OBs, whereas day 5 cells contained a large proportion of PPCs (Lin-/Cd34-) (Fig. 3D). EdU incorporation indicated that MPCs were less proliferative than PPCs at both days 0 and 5, though bone fracture enhanced proliferation in both populations (Fig. 3D). In addition, qRT-PCR analysis of cells sorted from day 5 callus verified that the Cd34+ cells highly expressed MPC markers including *Cd34, Ly6a, Cd248* and *Clec3b*, whereas CD34-cells more strongly expressed myofibroblast markers as well as *Inhba* (Fig. 3E). Lastly, immunohistochemistry on day 5 fractures from *Col2/Td* mice revealed that many Td+ cells were positive for both *α*SMA and activin A (Fig. 3F). Together, the data above provide further evidence for the occurrence of MPCs and PPCs within the evolving fracture callus and validate the myofibroblastic phenotype of PPCs characterized also by high *Inhba* expression.

### Activin A stimulates proliferation and differentiation in periosteal progenitors

The spatiotemporal links between PPC expansion and *Inhba* expression during fracture repair progression above indicate that activin A may directly promote progenitor cell proliferation and differentiation. To test this possibility, we isolated tibial periosteal mesenchymal progenitors, seeded them in standard maintenance monolayer culture and treated them with recombinant activin A (100 ng/ml) or with a neutralizing monoclonal antibody against mouse activin A (nActA.AB; 100 µg/ml). Cell number analysis on day 3 indicated that activin A treatment did stimulate cell proliferation, whereas treatment with nActA.AB had inhibited it (Fig. 4A).

**Fig. 4.**
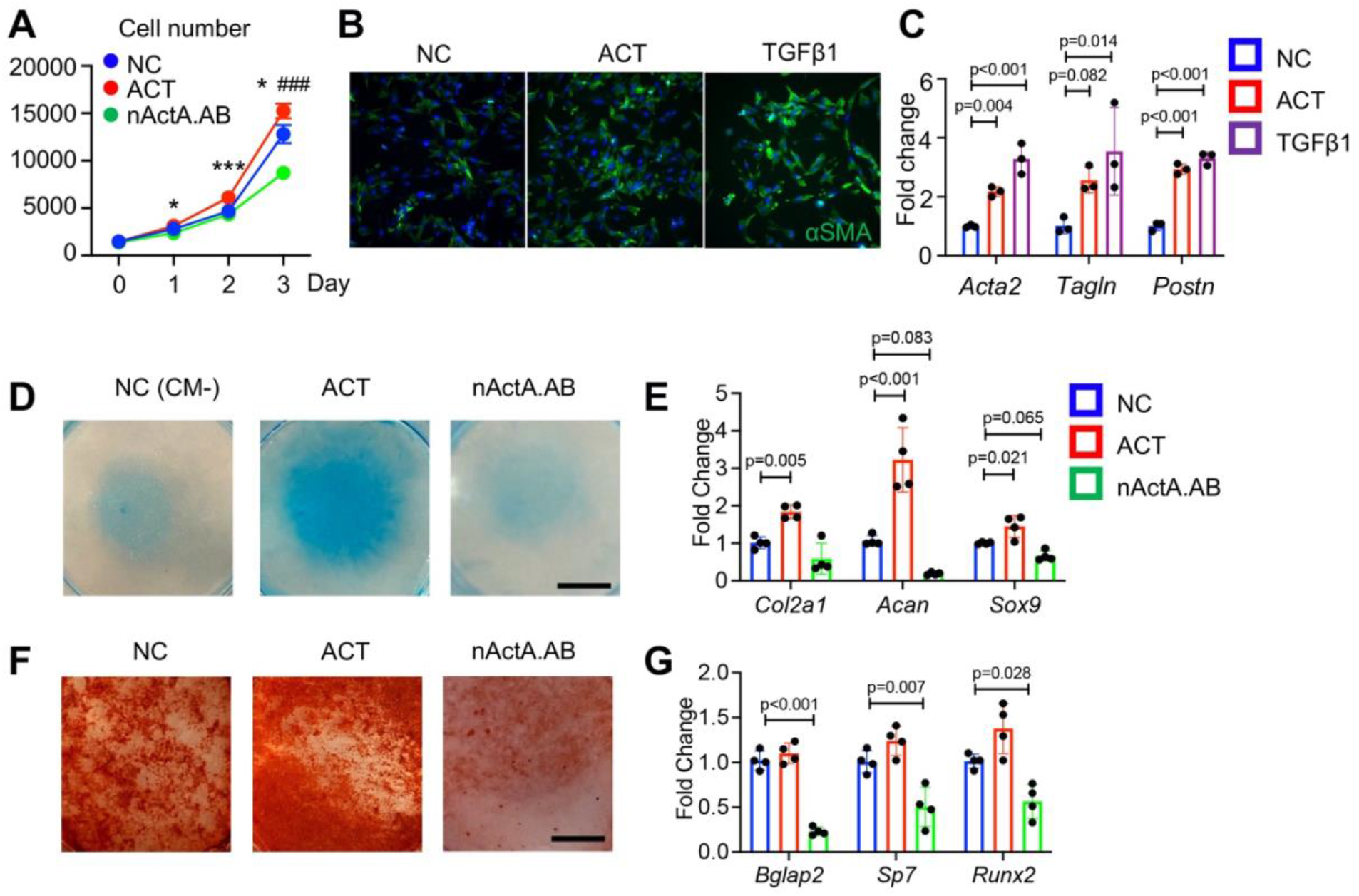
Activin A regulates proliferation and differentiation of periosteal progenitors in vitro. (**A**) Proliferation assay of periosteum mesenchymal progenitor cells treated with activin A (ACT) or neutralizing monoclonal antibody (nActA.AB). (**B**) Immunofluorescence images of αSMA in cultured periosteum mesenchymal progenitors treated with activin A or TGFβ1. (**C**) qRT-PCR data on expression of myofibroblast markers in periosteal mesenchymal progenitors after treatment with activin A or TGFβ1. (**D**) Alcian blue staining of periosteum mesenchymal progenitor cells in micromass undergoing chondrogenic differentiation in the presence of activin A or nActA.AB. (**E**) qRT-PCR analyses of chondrogenic markers on day 14 of culture in chondrogenic conditions. (**F**) Alizarin red staining of periosteum mesenchymal progenitor cells underging osteogenic differentiation in the presence of activin A or nActA.AB. (**G**) qRT-PCR analyses of osteogenic markers on day 14 of culture.

Remarkably, activin A treatment up-regulated the gene expression of myofibroblast markers including *Acta2* and *Tagln* (Fig. 4B and C) as did treatment with recombinant TGFβ1 which is known for its ability to promote myofibroblast development (*43*). Next, we tested whether activin A was able to stimulate chondrogenic and osteogenic cell differentiation that as predicted by our trajectories, is linked to the phenotypic bifurcation of PPCs (Fig. 2F). Thus, periosteal cell cultures were reared in basal chondrogenic or osteogenic media and were treated with activin A as above. Activin A treatment did stimulate chondrogenesis versus untreated cultures as revealed by strong alcian blue staining and higher expression of such cartilage markers as *Col2a1, Acan* and *Sox9* (Fig. 4, D and E). Activin A treatment did not appreciably enhance osteogenic differentiation (Fig. 4, F and G); nActA.AB treatment did, however, inhibit both osteogenesis and chondrogenesis (Fig. 4, D-G). Thus, endogenous and exogenous activin A acts to promote chondrogenic and osteogenic differentiation in periosteal progenitors.

### Systemic administration of activin A neutralizing antibody delays fracture repair

Given the apparent ability of activin A to stimulate periosteal progenitor proliferation and differentiation, it became reasonable to predict that the protein would have a positive and important role in fracture repair. To test this thesis, we utilized the same tibia fracture close model above, using 2 month-old WT mice. The animals were randomly divided into two groups. The first received biweekly injections of nActA.AB [immunoglobulin G2b (IgG2b) isotype at 10 mg/kg per injection] as in our recent HO study (*22*). The second group served as control and received injections of pre-immune IgG2b isotype antibody obtained from the same manufacturer, given at identical dose, route, and frequency. Based on the spatiotemporal patterns of fracture healing in this model (fig. S2), tibias from each group were harvested at 5, 7, 10 and 14 days from surgery to capture and analyze the cartilage and bone formation phases and at 6 weeks to measure the ultimate effectiveness of bone healing by mechanical tests. Histochemical analysis clearly showed that nActA.AB treatment significantly reduced overall callus size and cartilage and bone areas at all time points post fracture (5, 7, 10 and 14 days) compared to isotype antibody controls (Fig. 5, A and B). The changes in callus volume and bone volume were verified by μCT analysis (fig. S8). By 6 weeks in the control group, the tibial fractures were all bridged, indicating a successful recovery but those in the nActA.AB treatment group were lagging behind, leading to a decrease in fracture healing score (Fig. 5C). Furthermore, three-point bending analysis revealed 44.4%, 35.0%, and 29.0% reductions in energy to failure, stiffness and peak load, respectively, in injured tibias from nActA.AB-treated versus isotype-treated mice (Fig. 5D). To gain insights into whether the systemic nActA.AB treatment was affecting local cell populations and the PPCs in particular, we dissected out the fracture callus from nActA.AB-treated and isotype-treated mice on day 7 after fracture and processed the samples for gene expression analysis by qRT-PCR. We did find that nActA.AB treatment had decreased local endogenous expression of *Acta2, Sox9* as well as *Inhba* (Fig. 5E).

**Fig. 5.**
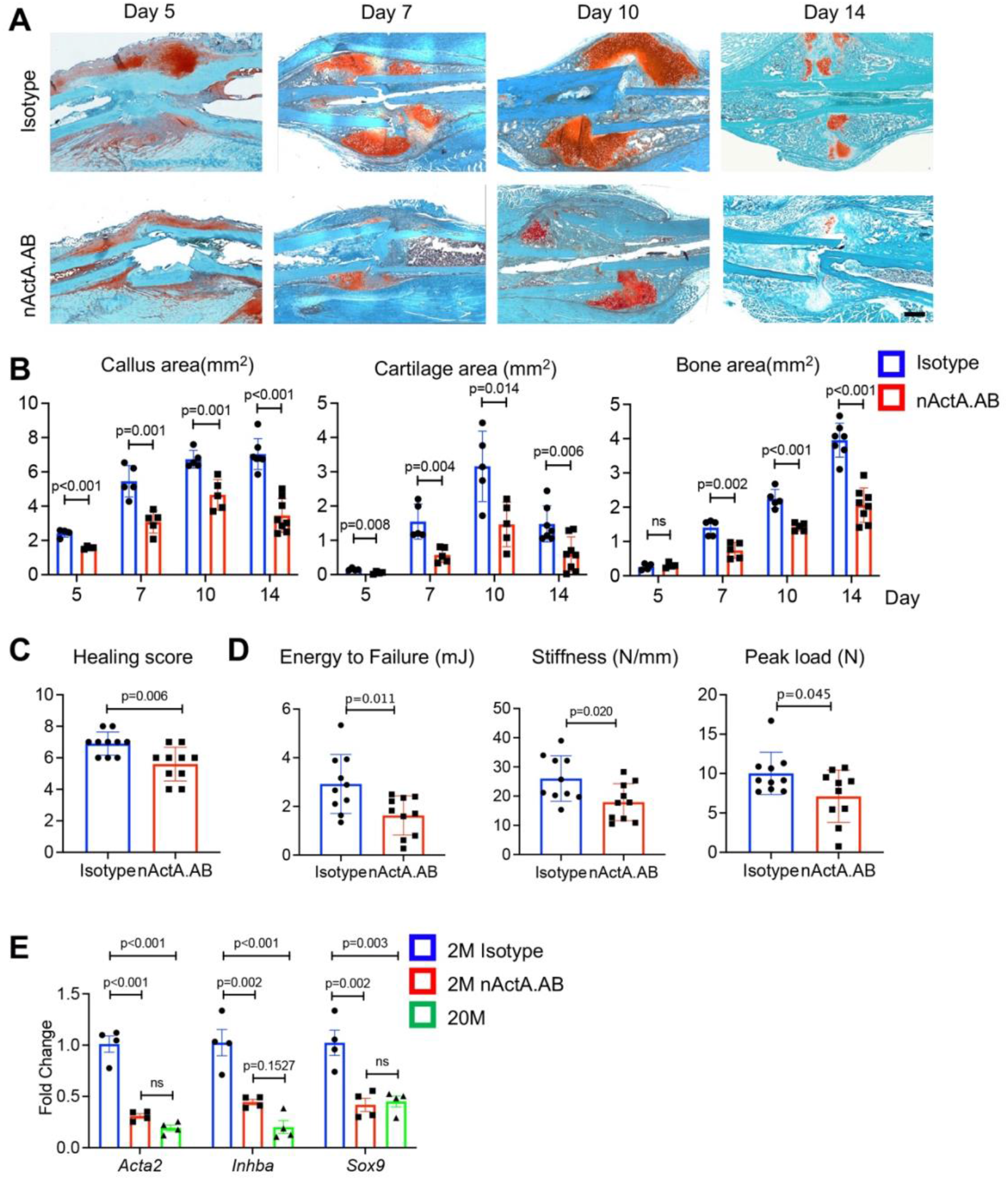
Systemic administration of activin A antibody delays mouse fracture healing. (**A**) Representative Safranin O/Fast green staining images of calluses at 5, 7, 10, and 14 days post fracture. Mice received injections of a control IgG2b isotype or a neutralizing monoclonal antibody against activin A (nActA.AB, 10 mg/kg) twice a week after fracture. Scale bar: 1000 μm. (**B**) Callus area, cartilage area, and bone area were measured at 5, 7, 10, and 14 days post fracture. n = 4-7 mice/time point. (**C**) Fracture healing scores were quantified at 6 weeks post fracture. n = 10 mice/treatment. (**D**) Four-point bending test was performed on bones at 6 weeks post fracture. n = 10 mice/treatment. (**E**) qRT-PCR analyses of *Acta2, Inhba*, and *Sox9* expression in day 7 callus from 2-month-old mice treated with nActA.AB and from 20-month-old mice without treatment. n = 4 mice/group.

### Local administration of activin A accelerates fracture healing

To complement the above studies, we carried out complementary gain-of-function studies and asked whether exogenous activin A may enhance fracture healing and represent a potential therapeutic. As above, we used a closed tibial fracture model with 2 month-old mice as well as older 20 month-old mice since older mice are more clinically relevant when testing a potential therapy. Immediately after fracture, a 50 μl aliquot of Matrigel containing recombinant activin A up to 1 μg was microinjected at the operated site; controls received Matrigel alone. Mice were then harvested at different time points to monitor the healing process. Notably, exogenous activin A implantation had increased callus size and cartilage and bone areas at each time point and in both age groups, based on histochemistry (Fig. 6, A and B) and μCT imaging (fig. S9).

**Fig. 6.**
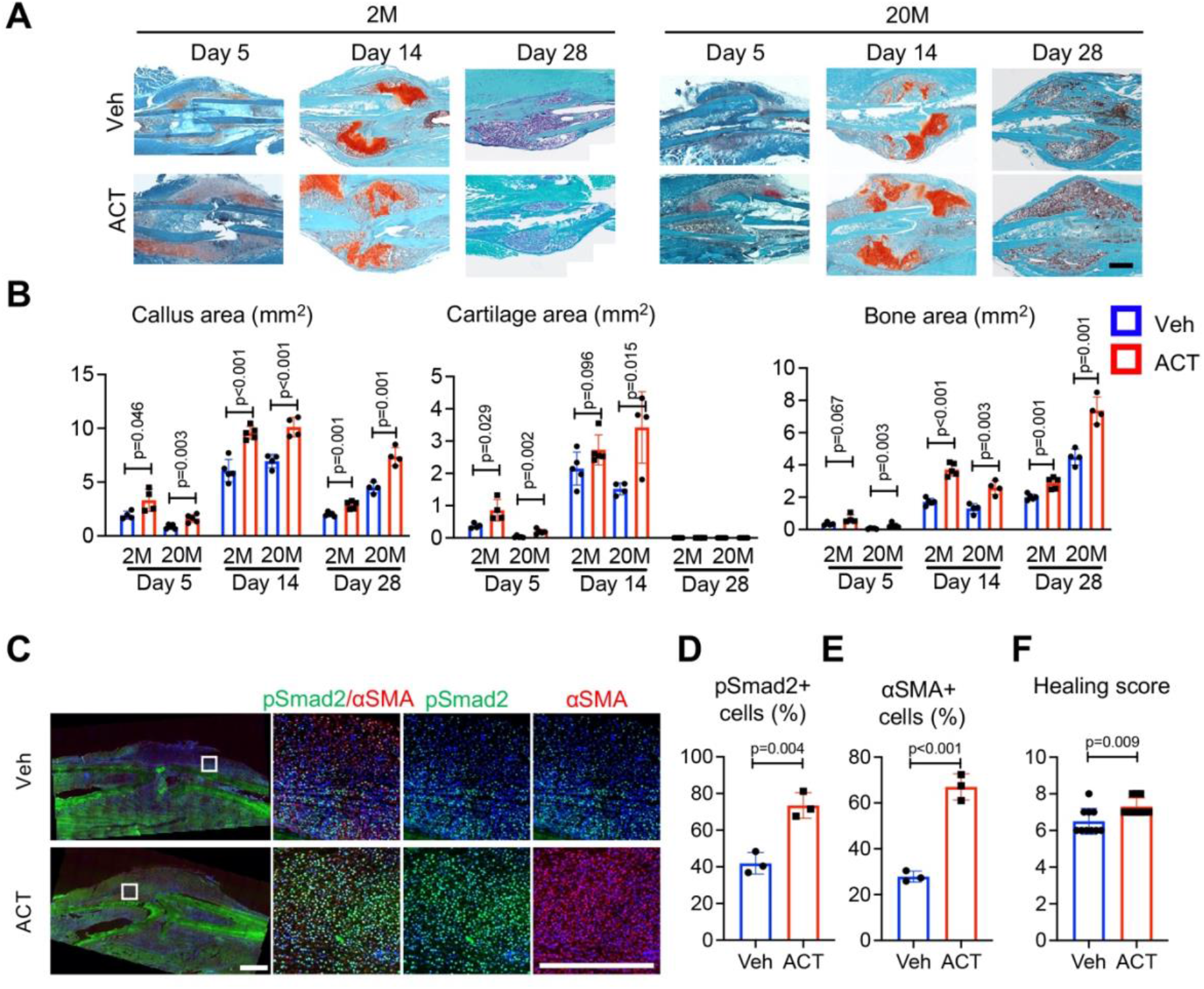
Local activin A implantation promotes fracture healing. (**A**) Representative Safranin O/Fast green staining images of fracture calluses at 5, 14 and 28 days post fracture. Mice at 2 or 20 months of age received a 50 μl Matrigel aliquot containing vehicle or activin A (1 μg) at the fracture site immediately after surgery. Scale bar: 1000 μm. (**B**) Callus area, cartilage area, and bone area were measured at 5, 14 and 28 days post fracture. n = 4 mice/group. (**C**) Immunofluorescence images of αSMA and pSmad2 in callus at day 5 post fracture. The boxed areas in the left images are shown enlarged on the right. (**D, E**) Quantified percentages of pSmad2+ cells (D) and αSMA+ cells (E) in callus. n = 3 mice/group. (**F**) Fracture healing scores quantified at 6 weeks post fracture. n = 10 mice/treatment.

Immunohistochemistry revealed that importantly, activin A implantation had elicited a major increase in the number of cells positive for pSMAD2 through which the protein normally elicits intracellular signaling and action (Fig. 6, C and D) (*44*). It was of interest to note also that the number of αSMA+ cells and the overall healing score were also increased by activin A implantation (Fig. 6, C, E and F), indicating an expansion of the resident PPC population and promotion of healing.

### Activin A promotes intramembranous bone defect repair

To strengthen our observations, we carried out additional loss- and gain-of-function experiments using a monocortical non-critical size (0.8 mm) drill-hole bone repair model that heals mainly through intramembranous ossification (*45*). Woven bone formation is usually observed by day 7 post-surgery, and bone bridging and re-corticalization occur by day 21 (*46*). Accordingly, drill-hole surgery was carried out mid-shaft in tibias of 2 month-old mice that were randomly divided into groups. For loss-of-function tests, mice were given biweekly injections of nActA.AB or isotype control as above. For gain-of-function tests, a 50 μl aliquot of Matrigel containing up to 1 μg of recombinant activin A was microinjected inside the medullary canal at the drill site; Matrigel alone was microinjected in companion controls. In all controls, bone formation became evident in the medullary region of interest by day 7 post-surgery and extensive bone formation in the drilled region had occurred by day 21 (Fig. 7A). Nearly all day 21 samples displayed complete defect bridging and based on μCT-based sagittal and cross-sectional reconstitution, volume fraction of reconstituted bone (BV/TV) was over 70% (Fig. 7B).

**Figure. 7.**
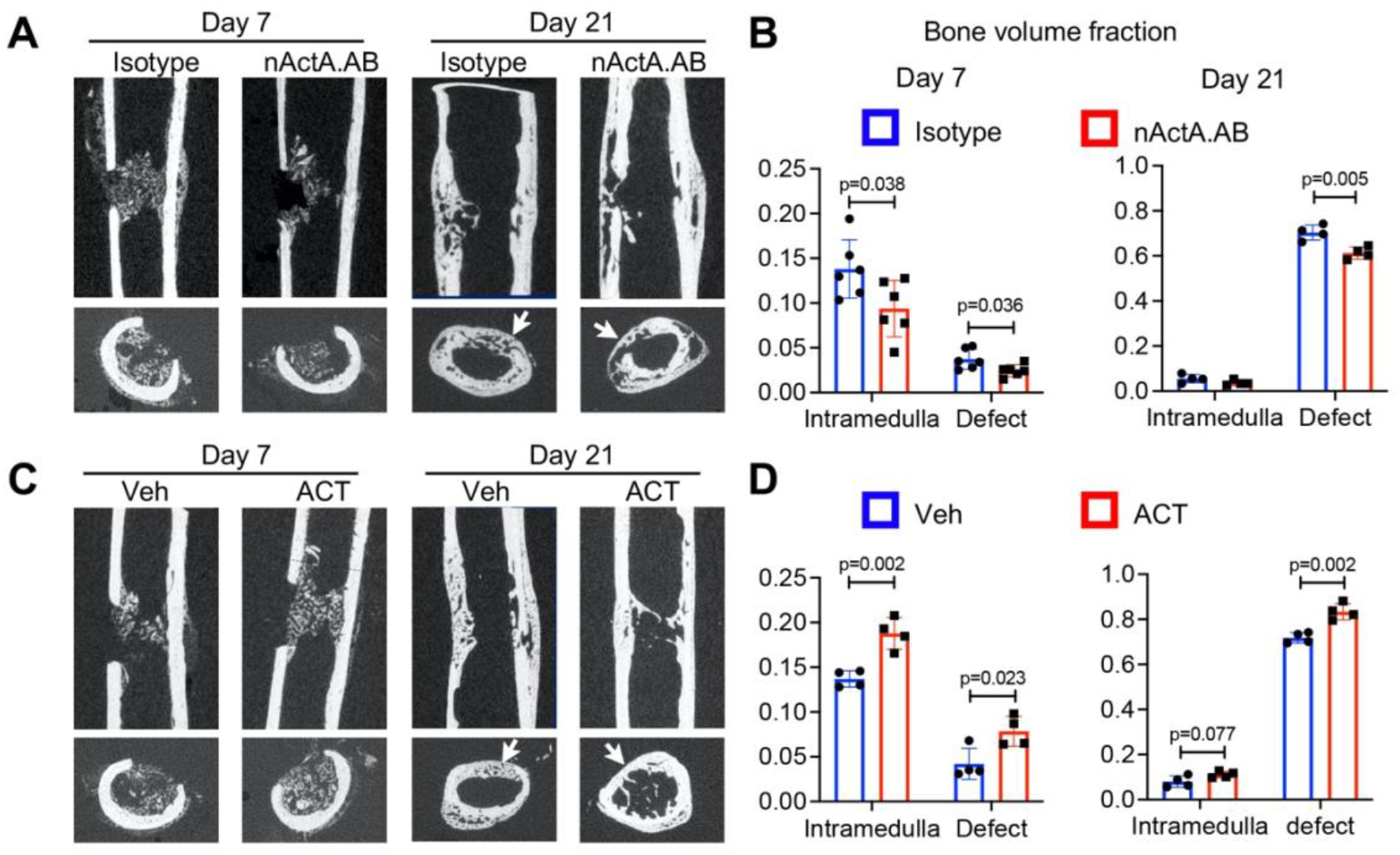
Activin A stimulates fracture healing in drill hole model. (**A**) Representative µCT images of mouse tibiae at 7 and 21 days post drill hole surgery. Mice received injections of a control IgG2b isotype or a neutralizing monoclonal antibody against activin A (nActA.AB, 10 mg/kg) twice a week after fracture. Top panel: bone longitudinal view; bottom panel, bone cross-sectional view. Arrows point to the original hole sites. (**B**) Bone volume fraction (BV/TV) of intramedullary area and defect area at 7 and 21 days post fracture. (**C**) Representative 2D µCT images of mouse tibiae at 7 and 21 days post drill hole injury. Mice received a 50 μl Matrigel aliquot containing vehicle or activin A (1 μg) at the drill site. Top panel: longitudinal view; bottom panel, cross-sectional view. Arrows point to the original hole sites. (**D**) Bone volume fraction (BV/TV) of intramedullary area and defect area at 7 and 21 days post fracture. n = 4-6 mice/time point.

Systemic nActA.AB administration caused an appreciable reduction in bone deposition in both drilled and medullary regions by day 7 and a significant drop by day 21 compared to isotype controls (Fig. 7A and B). Conversely, local microinjection of recombinant activin A significantly increased BV/TV in medullary and drilled regions at both day 7 and day 21 compared to vehicle controls (Fig. 7C and D).

## DISCUSSION

Our data identify a novel population of activin A-expressing proliferative progenitor cells (PPC) in the developing fracture callus and establish activin A as a potential new therapeutic tool to enhance fracture repair. We find that, in addition to its typical expression in inflammatory cells, activin A production and *Inhba* expression prominently mark the PPC population that rapidly expands within the enlarging callus. scRNAseq-based trajectory analysis indicates that the cells lie at the center of a developmental bifurcation eliciting the emergence of chondrocytes and osteoblasts within the repair mass (fig. S10). We demonstrate that endogenous activin A is required for the effective occurrence of these reparative processes and that exogenous activin A delivery enhances them, stimulating both intramembranous and endochondral ossification. We find also that exogenous activin A delivery robustly induces pSMAD2 signaling activity in vivo and promotes cell-autonomous progenitor differentiation down the myofibroblastic, chondrogenic and osteogenic lineages in vitro. In sum, this study demonstrates that activin A is an overall orchestrator and regulator of the fracture repair process, characterizes the prominent and previously unidentified PPC population, and represents a potential new therapeutic tool to promote bone healing.

Periosteal mesenchymal stem and progenitor cells are well known to be essential for fracture repair, but the sequential steps needed to turn them into reparative cells and mechanisms underlying this multifaceted progression have remained unclear (*1, 2*). The novel PPC population identified here occupies at an intermediate developmental step along the repair cascade and possesses a myofibroblast character (*47*). Prior to fracture injury, the periosteum contains very few PPCs, but fracture triggers a concerted response in which MPCs are rapidly activated to become PPCs that then differentiate into chondrocytes and osteoblasts. Based on cell prevalence, the PPCs appear to represent the bulk of cells within the thickening periosteum a few days after fracture. Notably, the PPCs express not only *Inhba* but also *Acta2* that encodes αSMA, the well-established marker of periosteal mesenchymal progenitors engaged in fracture callus development (*48*). Using *Acta2-CreER* mice, the latter research group recently demonstrated that αSMA+ cells constitute about 4% of Cd45-Ter119-Cd31- cells in homeostatic mouse periosteum and that DTA-mediated ablation of the cells severely reduces callus size after fracture (*7*). In the tissue injury and regeneration field, αSMA is widely used as a marker of myofibroblasts, a cell type first discovered in skin wound healing studies and then identified as a key player in many tissue repair processes (*47, 49*). A recent scRNA-seq study of fibroblast populations isolated from 13 injured and diseased mouse tissues including bone, and identified an Lrrc15+ cell cluster that displays a αSMA+ myofibroblast cell character and emerges only after injury (*50*). Interestingly, Lrrc15 is also a marker for PPCs. In addition, abundant αSMA+ stromal cells were recently reported to occur around the injury site after metal implant surgery in mouse tibiae (*51*). In sum, acute injury resulting from bone fracture, metal implantation or other insults appears to elicit a common and forceful repair response that is coupled to the emergence of progenitors with a myofibroblastic phenotype expressing genes including *Lrrc15, Acta2* and *Inhba*. Ongoing studies are directed toward deciphering more precisely what roles the myofibroblastic PPCs perform in bone repair, what function *Inhba* expression has during that stage, and what mechanisms ensure the controlled regression of the cells and involution of the callus once bone fracture repair is completed.

While these and other important issues remain to be tackled, our data provide compelling insights into how the repair process could be influenced therapeutically. As indicated above, activin A is a product of inflammatory cells, and initial interactions and cross talks between inflammatory cells and progenitor skeletogenic cells are recognized as essential for effective fracture repair (*2, 52, 53*). Inflammatory cells present at the fracture repair site include neutrophils and macrophages and release a spectrum of inflammatory and chemotactic mediators including members of the TNF and IL protein families (*1, 2*). These and other proteins are thought to lead to recruitment of fibroblasts, MSCs and skeletogenic progenitors from local sources, propelling the next phase of skeletal tissue repair production and deposition (*54, 55*), but details remain scant (*2*). The previously described roles of activin A in inflammation (*24*) and our current findings suggest that the initial inflammatory response needed for setting the repair process in motion may involve, and rely on, activin A itself. This possibility is further supported by our previous report showing that activin A is produced by skeletogenic cells (*22*). The present study builds on those findings and reveals more critically that *Inhba* becomes prominently expressed by the myofibroblastic PPCs within the callus and that activin A promotes chondrogenesis and osteogenesis in periosteal cells. Together, the data lead to the important notion that activin A could represent a regulatory and functional nexus linking inflammation to both local skeletogenic responses and fracture repair. Its capacity to play such diverse roles could endow the protein with ideal characteristics as a therapeutic for fracture repair deficiencies. A variety of means have been tested in both animal studies and patients to improve fracture healing, but an effective and safe therapy is yet to emerge and be clinically applied (*1, 56*).

Anabolic therapies utilize exogenous agents such as members of the BMP, FGF, Wnt or hedgehog protein families and may not be as effective as desired because of their targeting mainly a given step or a given population (*56*). Activin A may prove more effective because of its early presence from the very onset of the fracture repair process and its activity through sequential populations, including our newly described myofibroblastic node toward terminal differentiation of cartilage and bone cells. While this paradigm remains to be fully tested, the spectrum of biological action by activin A and the appreciable promotion of fracture repair we have demonstrated provide rational foundation and premise for the investigation of activin A for clinical application.

## MATERIALS AND METHODS

### Study design

The current study focused on testing and deciphering the roles of activin A in mouse long bone fracture repair. We first asked whether the protein was present at sites of experimental tibia fracture. We then characterized the cell populations recruited at the fracture site using *Col2Cre/Td* labeling and sorting followed by scRNAseq analyses, paying particular attention to their phenotypic differences and their developmental trajectories and evolution. Lastly, we directly tested the roles of activin A in fracture repair by performing loss-of-function and gain-of-function experiments, using a monoclonal neutralizing antibody and recombinant activin A, respectively. Samples sizes were determined based on prior experience and are indicated in figure legends. Mice were randomly assigned into different groups as needed.

### Animals

Specific pathogen-free 2-month-old *C57B16* female mice were purchased from the Jackson Laboratory. *Col2-Cre Rosa-tdTomato* (*Col2/Td*), and *Adipoq-Cre Rosa-tdTomato* (*Adipoq/Td*) mice were generated by breeding *Rosa-tdTomato* (Jackson Laboratory) mice with *Col2-Cre* (*57*) and *Adipoq-Cre* (*58*) mice respectively. Closed transverse fractures were made on right tibiae via a blunt guillotine with a pre-inserted intramedullary pin. For nActA.AB treatment, 10 mg/kg nActA.AB or pre-immune control IgG2b isotype were subcutaneously injected twice per week after fracture. For activin A treatment, growth factor-reduced, phenol red-free Matrigel (Corning) was mixed with activin A (1 μg) per 50 μl. To apply Matrigel mix, tibial anterior side was minimally exposed immediately after fracture and then, the Matrigel mix was applied immediately surrounding the fracture site using an insulin syringe. The skin was then closed with sutures. Bones were harvested at indicated times for histology and µCT analysis.

### Periosteum Td^+^ cell isolation and cell sorting

Periosteum cells were harvested as described previously (*30*). For day 0 before fracture and day 5 after fracture samples, mouse tibiae were dissected free of surrounding tissues and both ends were sealed with 3% agarose gel. The remaining bone fragments were digested in 2 mg/mL collagenase A and 2.5 mg/mL trypsin. Cells from the first 3 min of digestion were discarded and cells from a subsequent 30 min of digestion were collected as periosteal cells. For day 10 after fracture samples, mouse tibiae were dissected free of surrounding tissues, fracture callus were cut off using surgical blade and cut into small pieces. The callus fragments were digested in 2 mg/mL collagenase A and 2.5 mg/mL trypsin for 1 hour and collected as callus cells. For sorting, those cells were resuspended into FACS buffer containing 25 mM HEPES (Thermo fisher scientific) and 2% FBS in PBS and sorted for Td^+^ cells using Influx B or Aria B (BD Biosciences).

### Single-cell RNA sequencing of endosteal bone marrow cells

We constructed 3 batches of single cell libraries for sequencing: periosteum Td^+^ cells from day 0 before fracture (n=5 mice), day 5 after fracture (n=6 mice), day 10 after fracture (n=6 mice). 20,000 cells were loaded in aim of acquiring one single library of 10,000 cell for each time point by Chromium controller (V3 chemistry version, 10X Genomics Inc), barcoded and purified as described by the manufacturer, and sequenced using a 2×150 pair-end configuration on an Illumina Novaseq platform at a sequencing depth of ∼400 million reads. Cell ranger (Version 6.0.1, https://support.10xgenomics.com/single-cell-geneexpression/software/pipelines/latest/what-is-cell-ranger) was used to demultiplex reads, followed by extraction of cell barcode and unique molecular identifiers (UMIs). The cDNA insert was aligned to a modified reference mouse genome (mm10).

Seurat package V3 (*59*) was used for individual or integrated analysis of the datasets. Standard Seurat pipeline was used for filtering, variable gene selection, dimensionality reduction analysis and clustering. Doublets or cells with poor quality (genes>6000, genes<200, or >5% genes mapping to mitochondrial genome) were excluded. Expression was natural log transformed and normalized for scaling the sequencing depth to a total of 1×10^4^ molecules per cell. Seurat Cell-cycle scoring function were used to analyze cell proliferation, proliferative cells were defined as cells in G2M or S Phase. First identify the top 2000 variable genes by controlling for the relationship between average expression and dispersion. Then, expression matrix were scaled by regressing out cell cycle scores (G2M.Score and S.Score). Statistically significant principal components (PC) were selected as input for uniform manifold approximation and projection (UMAP) plots. For the integrated dataset, batch integration was performed using Harmony (version 1.0) (*60*). Different resolutions for clustering were used to demonstrate the robustness of clusters. In addition, differentially expressed genes within each cluster relative to the remaining clusters were identified using FindMarkers within Seurat. Sub-clustering was performed by isolating the mesenchymal lineage clusters using known marker genes, followed by reanalysis as described above. Gene ontology analysis and Gene Set Enrichment Analysis(GSEA) was performed using the clusterProfiler package (*61*).

To computationally delineate the developmental progression of periosteal mesenchymal cells and order them in pseudotime, we performed the trajectory analysis using (*62*). Briefly, UMAP was used as dimensional reduction after the PCA were calculated for individual or integrated datasets. Then Seurat objects were transformed into SingleCellExperiment objects. Slingshot trajectory analysis was conducted using the Seurat clustering information and with dimensionality reduction produced by UMAP.

RNA velocity analysis was performed as described. First, spliced and unspliced counts were generated using the Velocyto (*63*). Next, we used above-described Seurat pipeline to generate UMAP projection and clustering information. Finally spliced, unspliced matrix, clustering and UMAP projection information were input to Scvelo for visualizing the RNA (*42*).

### Micro-computed tomography (µCT) analysis

Tibiae harvested post fracture were scanned at the fracture sites by VivaCT 40 (Scanco Medical AG) at a 7.4 µm isotropic voxel size to acquire a total of 1000 µCT slices centering around the fracture site. A semi-automated contouring method was used to determine the callus perimeter. Briefly, callus and cortical bone sections were manually identified on the first slice and then had spline interpolation between points. The points were chosen on slices no more than ten slices apart (0.074 mm). These points were manually reviewed across every slice and re-interpolation was performed if necessary. All images were first smoothed by a Gaussian filter (sigma=1.2, support=2.0) and then applied by a threshold corresponding to 30% of the maximum available range of image gray scale values to distinguish mineralized tissue from unmineralized and poorly mineralized tissue. Callus region surrounding cortical bone was contoured for trabecular bone analysis. Based on µCT images, 6 weeks fracture samples were assigned fracture healing scores according to an 8-point radiographic scoring system(*64*). For drill hole model, the contouring of defect area or intramedullary area were manually defined in the similar way as fracture callus and followed by trabecular bone analysis.

### Mechanical testing

Tibiae harvested at 6 weeks after fracture were placed on a 4-point bending fixture and loaded with mechanical force at the previously fractured site using an Instron 5542 (Instron). The force to failure curve was recorded for analyzing peak load, stiffness, and energy to failure.

### Histology and immunohistochemistry (IHC)

Fractured tibiae were fixed in 4% PFA, decalcified in 10% EDTA for 3 weeks, and processed for paraffin embedding. A series of 6 μm-thick longitudinal sections were cut across the entire fracture callus from one side of cortical bone to the other side of cortical bone. For each bone, a central section with the largest callus area as well as two sections at 192 μm (∼1/4 bone width) before and after the central section were stained with Safranin-O/Fast green and quantified for cartilage area, bone area, and fibrosis area by ImageJ. After antigen retrieval, slides were incubated with primary antibodies, such as rabbit anti-pSmad2 (S465/467) (Cell Signaling, 3108S), mouse anti-αSMA (Sigma, A5228) at 4°C overnight, followed by Alexa Fluor 488 donkey anti-rabbit (Invitrogen, A-21246), Alexa Fluor 555 donkey anti-mouse (Invitrogen, A-31570) secondary antibodies. For EdU staining, mice received 1.6 mg/kg EdU at 3 h before sacrifice and the staining was carried out according to the manufacturer’s instructions (Click-iT™ Plus EdU Alexa Fluor™ 647 Flow Cytometry Assay Kit, Thermo Fisher Scientific; c10340).

To obtain whole mount sections for immunofluorescent imaging, freshly dissected bones were fixed in 4% PFA for 1 day, decalcified in 10% EDTA for 4-5 days, and then immersed into 20% sucrose and 2% polyvinylpyrrolidone (PVP) at 4°C overnight. Then sample was embedded into 8% gelatin in 20% sucrose and 2% PVP embedding medium. Samples were sectioned at 50 µm in thickness. Sections were incubated with primary antibodies, such as mouse anti-αSMA, goat anti-Activin A (R&D, AF338), at 4°C overnight, followed by Alexa Fluor 488 donkey anti-mouse, Alexa Fluor 647 donkey anti-goat (Invitrogen, A-21447) secondary antibodies incubation 1 hour at RT.

### Cell culture

Digested periosteal cells were seeded in the growth medium (αMEM supplemented with 15% fetal bovine serum (FBS) plus 55 μM β-mercaptoethanol, 2 mM glutamine, 100 IU/ml penicillin and 100 µg/ml streptomycin) for periosteal mesenchymal progenitor culture. For colony-forming unit fibroblast (CFU-F) assay, cells were seeded at 0.3×10^6^ cells/T25 flask. Seven days later, flasks were stained with 3% crystal violet to quantify CFU-F numbers. For cell proliferation assay, 1000 cells were seeded into 96 well plate in growth medium containing following reagents Act A (100 ng/ml) or nActA.AB (100 μg/ml). Cell numbers at day 0, 1, 2 and 3 were quantified using CyQUANT Proliferation Assay Kit (Invitrogen, C35011); For myofibroblast differentiation assay, 0.2 ×10^6^ cells were seeded into 6 well plate with serum-free medium containing Act A (100 ng/ml) or TGF-β1 (10 ng/ml) for 72 hrs, followed by staining with mouse anti-αSMA primary antibody for 1 hr and Alexa Fluor 488 donkey anti-mouse secondary antibody for 1 hr at room temperature. For osteogenic differentiation assay, when reach confluent, cells were switched to osteogenic medium (αMEM with 10% FBS, 10 Nm dexamethasone, 10 mM β-glycerophosphate, 50 μg/mL ascorbic acid, 100 IU/ml penicillin and 100 µg/ml streptomycin) containing following reagents Act A (100 ng/ml) or nActA.AB (100 μg/ml) for 2 weeks followed by alizarin staining. For chondrogenic differentiation, micromass cultures were initiated by spotting 20 μl of the cell suspension (0.5 × 10^6^ cells/spot) onto the surface of 24-well tissue culture plates. After a 2 hr incubation at 37°C in a humidified CO^2^ incubator to allow for cell attachment, the cultures were switched to basic chondrogenic medium (high glucose DMEM, 100 µg/ml sodium pyruvate, 1% ITS+ Premix, 50 µg/ml ascorbate-2-phosphate, 40 μg/ml L-proline, 0.1 mM dexamethasone, 100 IU/ml penicillin and 100 µg/ml streptomycin) containing following reagents Act A (100 ng/ml); nActA.AB (100μg/ml) for two weeks followed by alcian blue staining.

### qRT-PCR analysis

Sorted cells or cultured cells were collected in TRIzol Reagent (Sigma, St. Louis, MO, USA). A Taqman Reverse Transcription Kit (Applied BioSystems, Inc., Foster City, CA, USA) was used to reverse transcribe mRNA into cDNA. Following this, quantitative realtime PCR (qRT-PCR) was performed using a Power SYBR Green PCR Master Mix Kit (Applied BioSystems, Inc). The primer sequences for the genes used in this study are listed in Supplementary Table 1.

### Study approval

The experimental protocols were approved by the Institutional Animal Care and Use Committees of the University of Pennsylvania and the Children’s Hospital of Philadelphia. The experiments were performed in the animal facilities of our Institutions that implement strict regimens for animal care and use. In accordance with the standards for animal housing, mice were group housed at 23-25°C with a 12 hr light/dark cycle and allowed free access to water and standard laboratory pellets.

### Statistical methodologies

Data are expressed as means ± standard deviation (SD) and analyzed by t-tests or one-way ANOVA with Tukey posttest for multiple comparisons using Prism software (GraphPad Software, San Diego, CA). For cell culture experiments, observations were repeated independently at least three times with a similar conclusion, and only data from a representative experiment are presented. Values of p<0.05 were considered significant.

## Supplementary Materials

Fig. S1. *Inhba* expression in mouse bone marrow.

Fig. S2. *Col2/Td* labels periosteal mesenchymal progenitors in intact and fractured tibiae.

Fig. S3. Large scale scRNA-seq analysis of Td+ cells sorted from tibial periosteum of 2 month-old *Col2/Td* mice.

Fig. S4. PPCs are greatly expanded after fracture.

Fig. S5. The expression patterns of previously reported periosteal mesenchymal progenitor markers.

Fig. S6. Myofibroblast marker genes expression pattern during fracture healing process.

Fig. S7. RNA velocity analysis indicates the differentiation route of periosteal mesenchymal lineage cells during fracture healing.

Fig. S8. Blocking activin A activity impedes mouse fracture healing.

Fig. S9. Activin A treatment accelerates fracture healing.

Fig. S10. Graphic abstract

## Acknowledgments

We thank Dr. Ivo Kalajzic at the University of Connecticut Health Center for critical reading of the manuscript and suggestions, and the Penn Center for Musculoskeletal Disorders (PCMD) at the University of Pennsylvania for expert use of its histology, µCT imaging and biomechanics core facilities. The PCMD is supported by the NIH grant P30AR069619.

## Author contributions

MP and LQ designed the study. LY, LZ, YW, TG, LW, DR and CM carried out the animal experiments and imaging. LY and LZ performed in vitro experiments. LY, CM and LQ carried out the scRNA-seq data analyses. JA and JB provided technical and material support and consultation. LQ, LY and MP wrote the original manuscript draft. All authors participated in reviewing and editing. LQ and MP secured funding and supervised the project.

## Data and materials availability

Sequencing data have been deposited in GEO under accession code GSE192630 (reviewer password: yxojgggwnlutbgv). All other data needed to evaluate the conclusions of the paper are present in the paper or the Supplementary Materials.

