## Supplementary material for "Activin A promotes bone fracture repair and acts through a novel myofibroblastic cell population in callus"

Fig. S1. *Inhba* expression in mouse bone marrow.

Fig. S2. *Col2/Td* labels periosteal mesenchymal progenitors in intact and fractured tibiae.

Fig. S3. Large scale scRNA-seq analysis of Td+ cells sorted from tibial periosteum of 2 month-old *Col2/Td* mice.

Fig. S4. PPCs are greatly expanded after fracture.

Fig. S5. The expression patterns of previously reported periosteal mesenchymal progenitor markers.

Fig. S6. RNA velocity analysis indicates the differentiation route of periosteal mesenchymal lineage cells during fracture healing.

Fig. S7. Myofibroblast marker genes expression pattern during fracture healing process.

Fig. S8. Blocking activin A activity impedes mouse fracture healing.

Fig. S9. Activin A treatment accelerates fracture healing.

Fig. S10. Graphic abstract.

Table S1: RT-PCR primer sequences

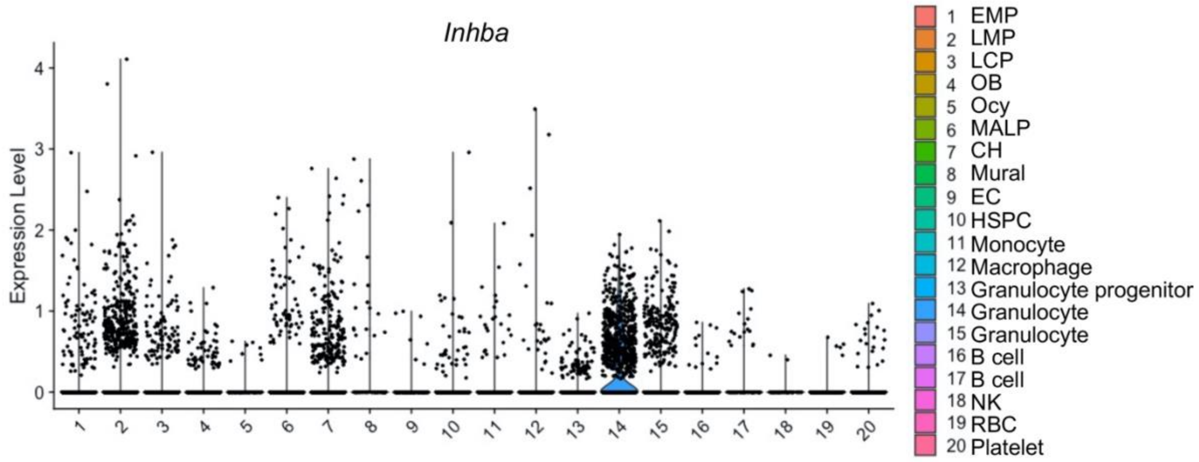


**Fig. S1. *Inhba* expression in mouse bone marrow.**

Violin plot of *Inhba* expression in bone marrow cells from 1-month-old mice based on our previous published scRNA-seq data.

EMP: early mesenchymal progenitor, LMP: late mesenchymal progenitor, LCP: lineage committed progenitor, OB: osteoblast, Ocy: osteocyte, MALP: marrow adipose lineage precursor, CH: chondrocyte, EC: endothelial cell, HSPC: haemopoietic stem and progenitor cell, NK: nature killer cell, RBC: red blood cell.

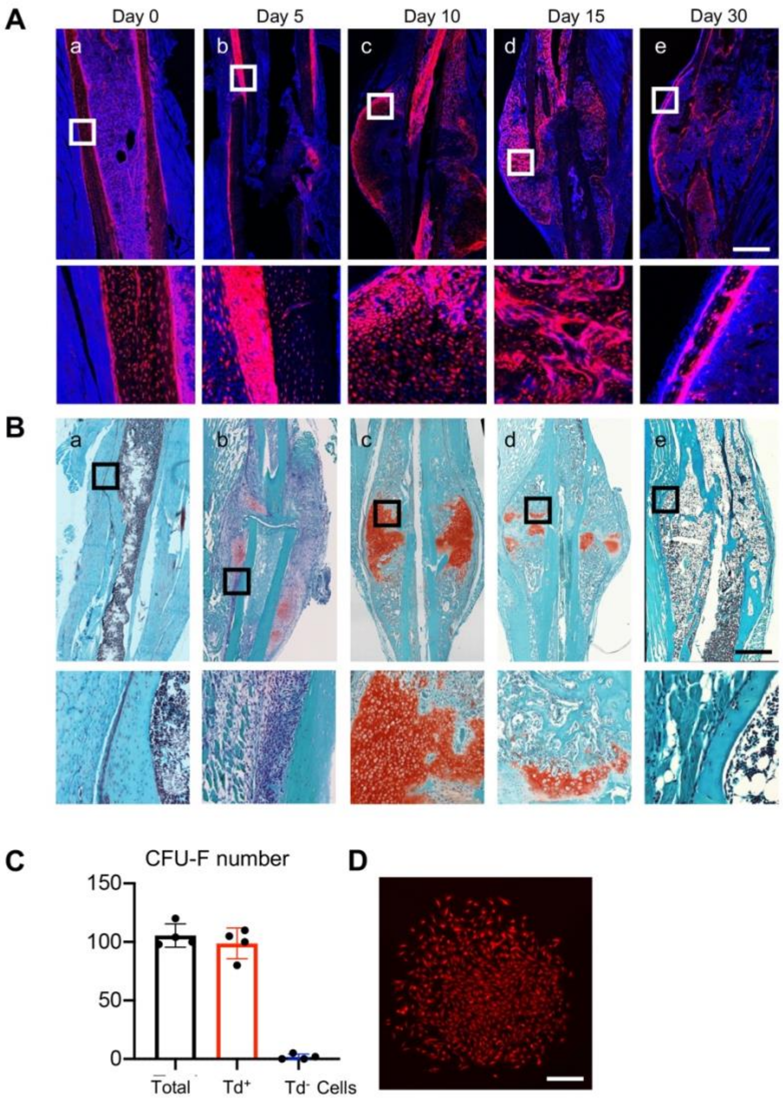


**Fig. S2. *Col2/Td* labels periosteal mesenchymal progenitors in intact and fractured tibiae.** (**A**) Fluorescence images of intact (day 0) and fractured tibiae from 2-month-old *Col2/Td* mice. Fractured samples were collected at days 5, 10, 15, and 30 post injury. The squared areas in the top panels are enlarged at the bottom panels. Scale bar: 1000 µ m. (**B**) Safranin-O staining of intact and fractured tibiae. The squared areas in the top panels are enlarged at the bottom panels. Scale bar: 1000µm. (**C**) Unsorted and sorted periosteal cells were cultured for CFU-F assay. Periosteal cells were digested from intact *Col2/Td* mice and sorted for Td+ cells and Td- cells. $1x10^6$ total digested cells, $1x10^4$ Td+ cells, and $1x10^6$ Td- cells were seeded per flask to count

CFU-F number 10 day later. n = 4 flasks/group. (**D**) Only Td+ cells formed CFU-F colonies.

Scale bar: 200 μm.

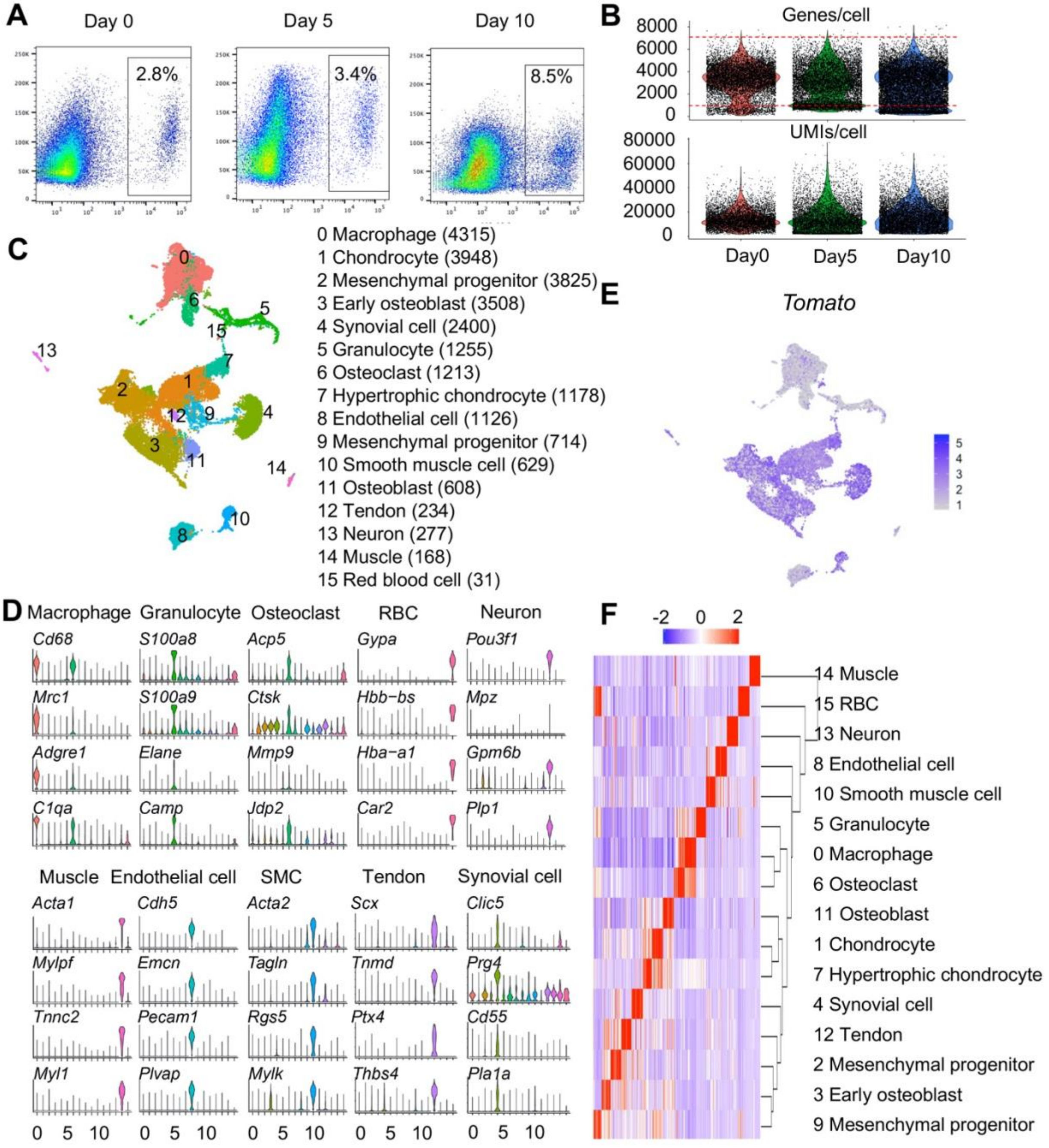


**Fig. S3. Large scale scRNA-seq analysis of Td+ cells sorted from tibial periosteum of 2 month-old *Col2/Td* mice**.

(**A**) Flow assay of digested periosteal cells from 2 month-old *Col2/Td* mice with or without fracture. Periosteal cells from intact tibiae (day 0) and injured tibiae at day 5 and day 10 post fracture were collected to measure the percentage of Td+ cells. n= 5-6 mice/time point. (**B**) Violin plots show average number of genes and UMIs per cell in a merged scRNA-seq dataset containing all sequenced cells at 3 time points (day 0, 5, and 10). Red box indicates cells within the selection criteria of quality controls. (**C**) The UMAP plot of 25429 Td+ periosteal cells in the merged scRNA-seq dataset (n = 5-6 mice/time point). Cell numbers are listed in parenthesis next to cluster names. (**D**) Violin plots of cluster-specific makers of hematopoietic cells ((macrophages, granulocytes (granulo), osteoclasts (OC) and red blood cells (RBC)), muscle cells, neuron cells, endothelial cells (ECs), smooth muscle cells (SMCs), tendon cells, and synovial cells. (**E**) The expression pattern of *Tomato* in the UMAP plot. (**F**) Hierarchy clustering and heatmap of all cell clusters. Color bar on the top indicates the gene expression level.

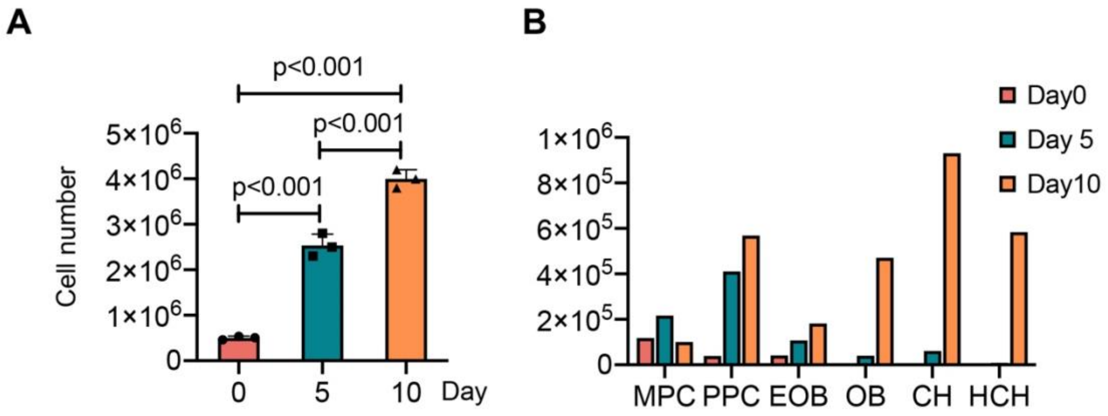


**Figure S4. PPCs are greatly expanded after fracture.**

(**A**) Total digested periosteal cells per bone were counted from intact (day 0) and injured (day 5 and 10 post fracture) mice at 2 months of age. n = 3 mice/time point. (**B**) Based on this information, the number of cells in each cell cluster at different time points was estimated. MPC: mesenchymal progenitor cell; PPC: proliferative progenitor cell; CH: chondrocyte; HCH: hypertrophic chondrocyte; EOB: early osteoblast; OB: osteoblast.

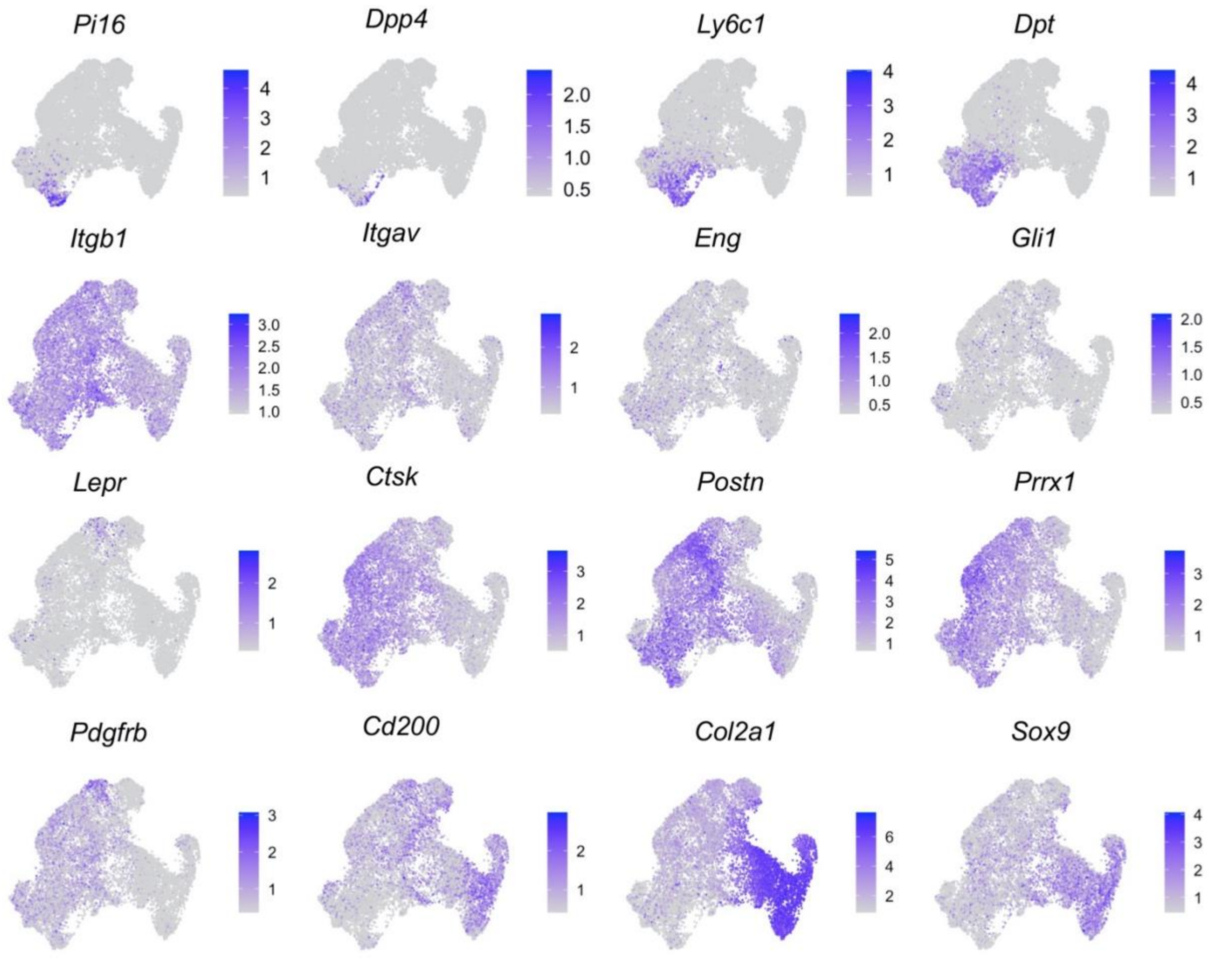


**Fig. S5. The expression patterns of previously reported periosteal mesenchymal progenitor markers.**

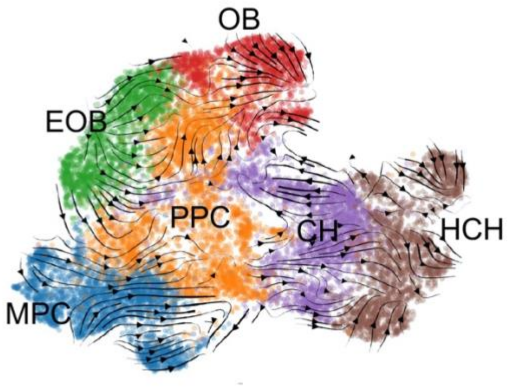


**Fig. S6. RNA velocity analysis indicates the differentiation routes of periosteal mesenchymal lineage cells during fracture healing.**

MPC: mesenchymal progenitor cell; PPC: proliferative progenitor cell; CH: chondrocyte; HCH: hypertrophic chondrocyte; EOB: early osteoblast; OB: osteoblast.

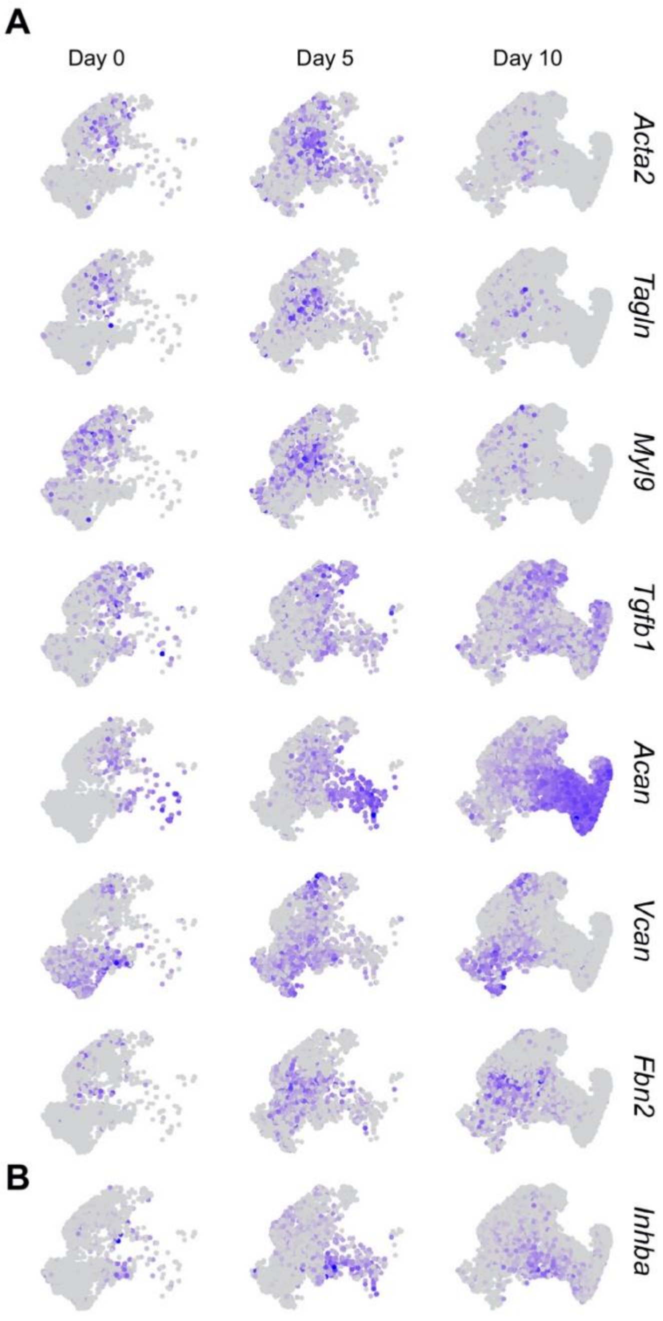


**Fig. S7. Myofibroblast marker genes expression pattern during fracture healing process.** **(A)** Myofibroblast marker genes expression pattern during fracture healing process. (B) *Inhba* gene expression pattern during fracture healing process.

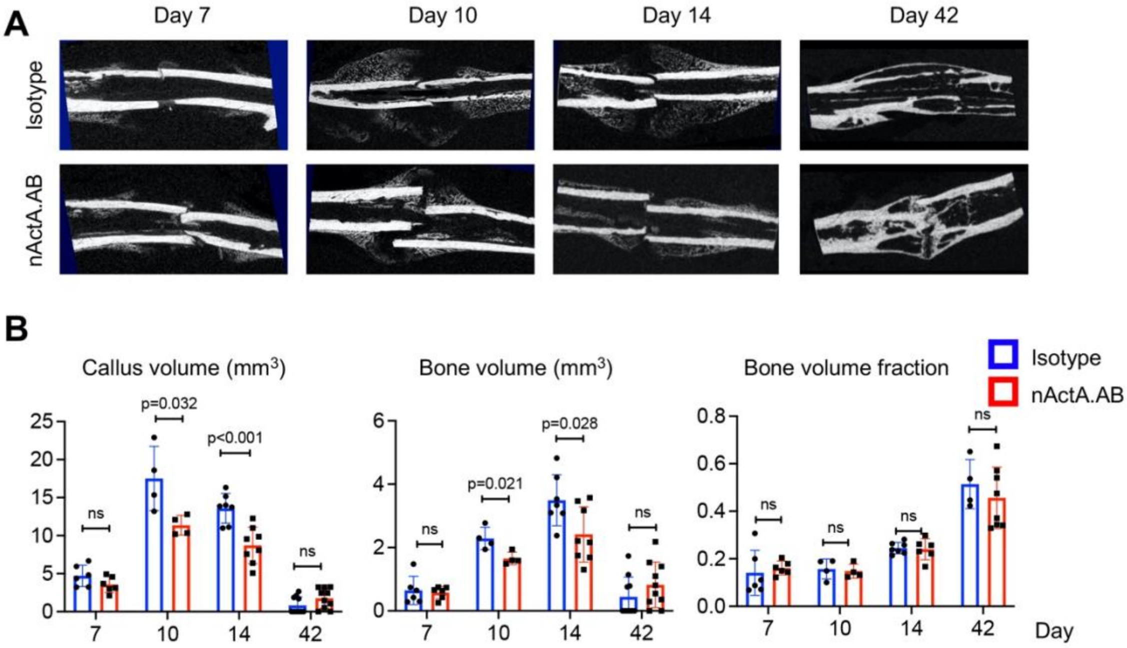


**Fig. S8. Blocking activin A activity impedes mouse fracture healing.**

(**A**) Representative μCT images of fracture calluses at 7, 10, 14, and 42 days post fracture. Mice received injections of an IgG2b isotype or a neutralizing monoclonal antibody against activin A (nActA.AB, 10 mg/kg) twice a week after fracture. (**B**) Callus volume, bone volume, and bone volume fraction of fracture calluses at 7, 10, 14, and 42 days post fracture were measured. n= 4-10 mice/treatment.

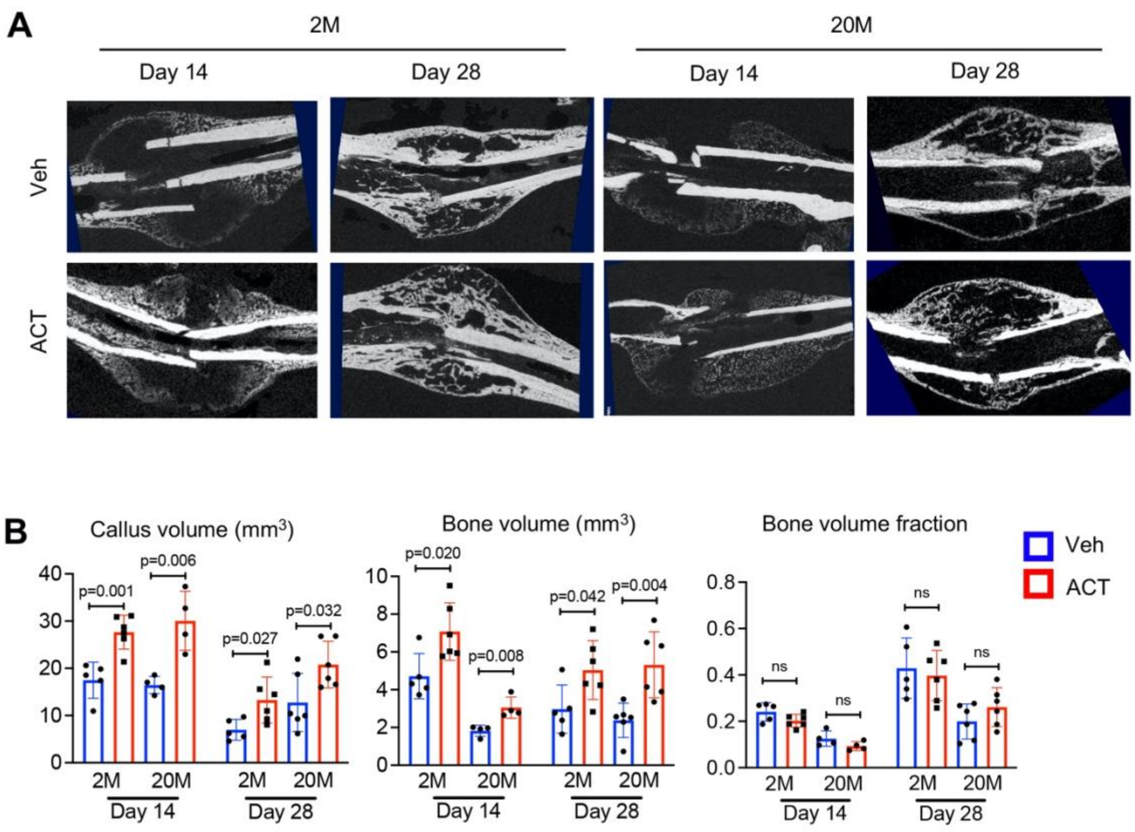


**Fig. S9. Activin A treatment accelerates fracture healing.**

(**A**) Representative μCT images of fracture calluses at 14 and 28 days post fracture. Mice at 2 or 20 months of age received 50 μl Matrigel containing vehicle or Activin A (1 μg) at the fracture site when fracture was made. (**B**) Callus volume, bone volume, and bone volume fraction of fracture calluses at 14 and 28 days post fracture were measured. n= 4-6 mice/group.

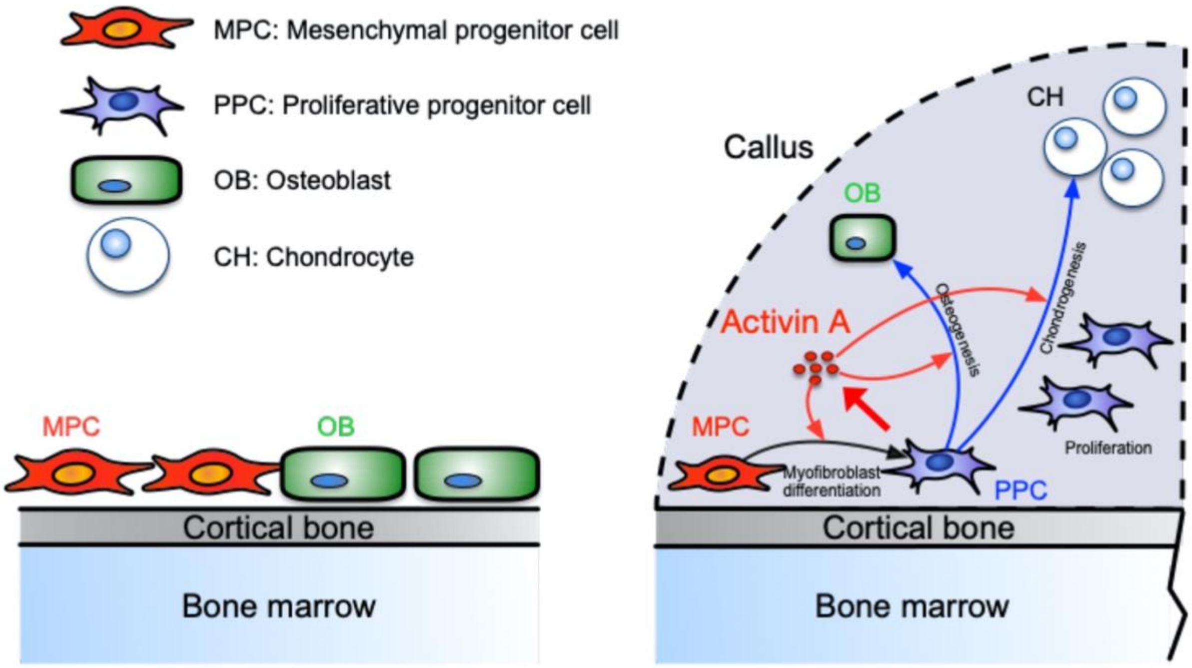


**Fig. S10. Graphic abstract depicting the proposed roles of activin A within the fracture callus.**

Table S1. Mouse real time RT-PCR primer sequences used in this study

| Gene | Forward primer | Reverse primer |
|---|---|---|
| *Cd34* | 5’- CTGGGTAGCTCTCTGCCTGAT -3’ | 5’- TGGTAGGAACTGATGGGGATATT-3’ |
| *Ly6a* | 5’-GAGTGGGAACTGGTAGTGTTG-3’ | 5’-CGCACAGAGCGATGAAGGT-3’ |
| *Cd248* | 5’-CGAGCCTCCTACTTCCAGTG-3’ | 5’-GGACAGGTAGCGATCCAGGT-3’ |
| *Clec3b* | 5’-CTGAACCGCTTTGGCAAGAC-3’ | 5’-GCCCTCTCTTATCGCCAGAT-3’ |
| *Acta2* | 5’-GTCCCAGACATCAGGGAGTAA-3’ | 5’-TCGGATACTTCAGCGTCAGGA-3’ |
| *Tagln* | 5’-CAACAAGGGTCCATCCTACGG-3’ | 5’-ATCTGGGCGGCCTACATCA-3’ |
| *Postn* | 5’-CCTGCCCTTATATGCTCTGCT-3’ | 5’-AAACATGGTCAATAGGCATCACT-3’ |
| *Col2a1* | 5’-GGGAATGTCCTCTGCGATGAC-3’ | 5’-GAAGGGGATCTCGGGGTTG-3’ |
| *Acan* | 5’-CCTGCTACTTCATCGACCCC-3’ | 5’-AGATGCTGTTGACTCGAACCT-3’ |
| *Sox9* | 5’-GAGCCGGATCTGAAGAGGGA-3’ | 5’-GCTTGACGTGTGGCTTGTTC-3’ |
| *Bglap2* | 5’-CTCTGTCTCTCTGACCTCAC-3’ | 5’-AGTCCTCTTAATCCTCGTGGG-3’ |
| *Sp7* | 5’-AGAGGTTCACTCGCTCTGACGA-3’ | 5’-TTGCTCAAGTGGTCGCTTCTG-3’ |
| *Runx2* | 5’-TAAAGTGACGGACGGTCCC-3’ | 5’-TGCGCCCTAAATCACTGAGG-3’ |
| *Inhba* | 5’-TGAGAGGATTTCTGTTGGCAAG-3’ | 5’-TGACATCGGGTCTCTTCTTCA-3’ |
| *Actb* | 5’-GGCTGTATTCCCCTCCATCG-3’ | 5’-CCAGTTGGTAACAATGCCATGT-3’ |
